# A single oscillating proto-hypothalamic neuron gates taxis behavior in the primitive chordate *Ciona*

**DOI:** 10.1101/2023.04.24.538092

**Authors:** Janeva Chung, Erin Newman-Smith, Matthew J. Kourakis, Yishen Miao, Cezar Borba, Juan Medina, Tao Laurent, Benjamin Gallean, Emmanuel Faure, William C Smith

## Abstract

*Ciona* larvae display a number of behaviors, including negative phototaxis. In negative phototaxis, the larvae first perform short spontaneous rhythmic casting swims. As larvae cast in a light field, their photoreceptors are directionally shaded by an associated pigment cell, providing a phototactic cue. This then evokes an extended negative taxis swim. We report here that the larval forebrain of *Ciona* has a previously uncharacterized single slow-oscillating inhibitory neuron (neuron *cor-assBVIN78*) that projects to the midbrain, where it targets key interneurons of the phototaxis circuit known as the *photoreceptor relay neurons*. The anatomical location, gene expression and oscillation of cor-assBVIN78 suggest homology to oscillating neurons of the vertebrate hypothalamus. Ablation of cor-assBVIN78 results in larvae showing extended phototaxis-like swims, but which occur in the absence of phototactic cues. These results indicate that cor-assBVIN78 has a gating activity on phototaxis by projecting temporally-oscillating inhibition to the photoreceptor relay neurons. However, in intact larvae the frequency of cor-assBVIN78 oscillation does not match that of the rhythmic spontaneous swims, indicating that the troughs in oscillations do not themselves initiate swims, but rather that cor-assBVIN78 may modulate the phototaxis circuit by filtering out low level inputs while restricting them temporally to the troughs in inhibition.

## INTRODUCTION

The tunicates, owing to their position in phylogeny as chordates and the closest extant relatives of the vertebrates (*1*), are valuable model animals for genetics, genomics, embryology and neurobiology. Of particular interest to researchers are the tadpole larvae of ascidians, such as those of the solitary ascidian *Ciona*. The central nervous system (CNS) of *Ciona* larvae highlights the common body plan of chordates: not only does it develop from a neural plate, but it also has an anterior-to-posterior anatomy that parallels that of vertebrates. Regions within the *Ciona* larva CNS show gene expression, neuron classification, and synaptic connectivity that tie them to vertebrate forebrain, midbrain, hindbrain and spinal cord [Fig. 1A; (*2*–*4*)]. For historical reasons these domains were called the *anterior sensory vesicle, posterior sensory vesicle, motor ganglion* and *caudal nerve cord*, respectively. Because of compelling data linking these CNS domains in *Ciona* and vertebrates, and for the sake of clarity, we will refer to them here by their more familiar vertebrate homolog names. Despite these similarities to vertebrate CNSs, the split of the tunicates and vertebrates is ancient, and, moreover, the *Ciona* larval CNS stands apart in its simplicity, having only ∼180 neurons.

**Figure. 1.**
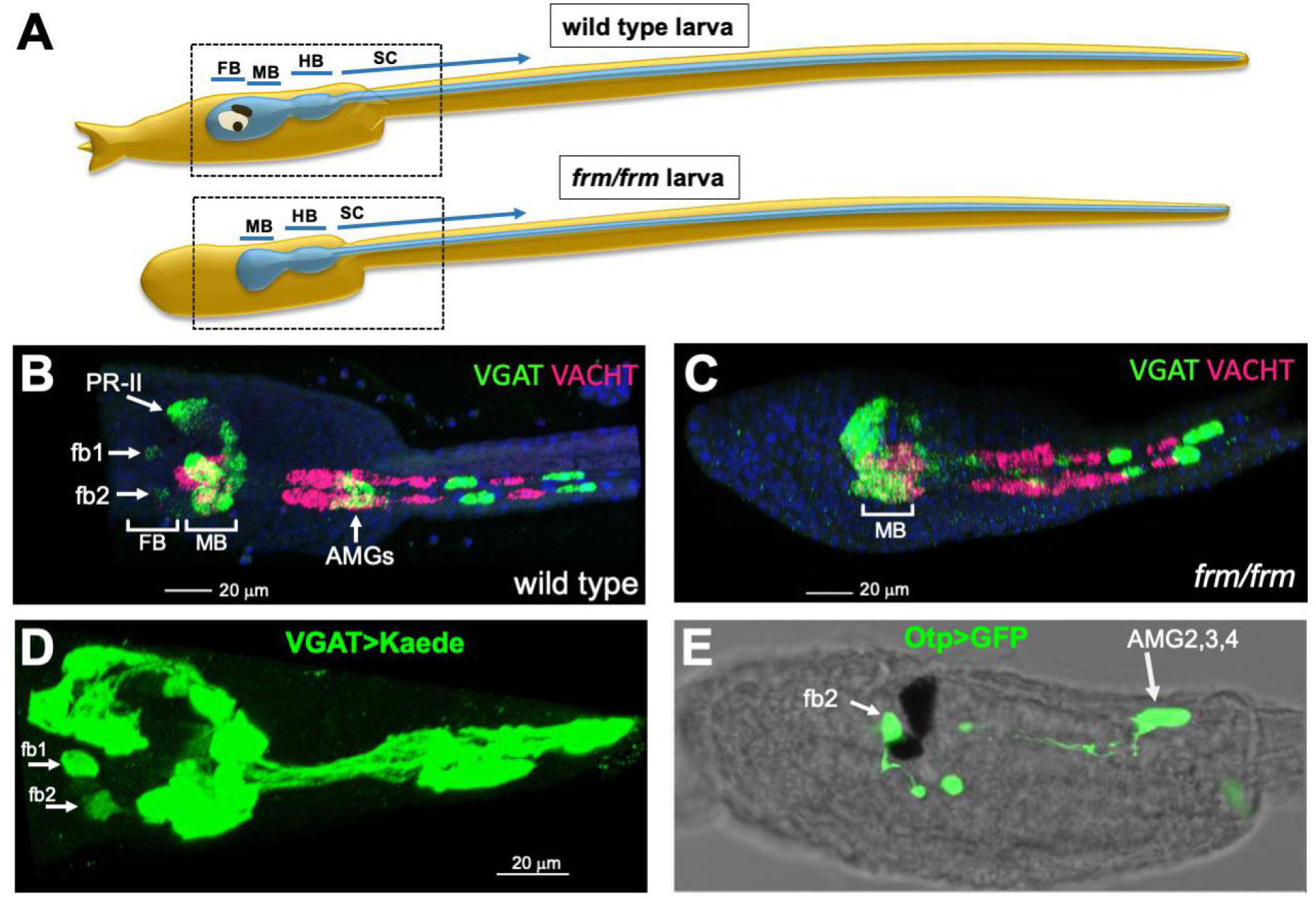
Inhibitory neurons in the *Ciona* larval forebrain. **(A)** Cartoons of wild type and *frm/frm* larvae. The CNS is depicted in blue and labeled with the major domains: forebrain (FB), midbrain (MB), hindbrain (HB), and spinal cord (SC). Top and bottom dashed boxes indicate the approximate regions shown in panels B and C, respectively. **(B and C)** Hybridization chain reaction *in situ* hybridization for VGAT and VACHT in wild type and *frm/frm* larvae, respectively. The two VGAT^+^ neurons labeled **fb1** and **fb2** are absent in the *frm/frm* mutant, as are the Group II photoreceptors (PR-II). **(D)** Expression of the fluorescent protein Kaede under control of the VGAT promoter. The image is a confocal projection of a larva immunostained with anti-Kaede antibodies. **(E)** fb2 and AMG2,3, and 4 are labeled with a reporter construct expressing GFP under control of the Otp promoter. Immunostaining with anti-GFP antibodies is shown. Anterior is to the left in all panels. See also Figure S1.

The availability of the *Ciona* larval connectome dataset, combined with easily assayed and quantified behaviors, has allowed for the investigation of individual neurons in circuits driving behaviors (*3, 5*–*7*). One such *Ciona* larval behavior is negative phototaxis. The minimal neural circuit for phototaxis is well described, and consists of a relay of four neuron types starting with photoreceptors to primary and then secondary interneurons, and finally to motor neurons (*6, 8*). Negative phototaxis is a robust behavior that appears in late-stage larvae, and is thought to aid in dispersal and settling (*9, 10*). Experimentally, *Ciona* larvae display negative phototaxis when presented with constant, but directional, illumination (Movie S1). Phototaxis initiates spontaneously with casting swims, in which larvae attempt to discern the direction of illumination with the aid of a pigmented cell that directionally shades the outer segments of their photoreceptors (*10*). As larvae turn away from the light, the pigment cell casts a shadow on the photoreceptors, providing a directional cue. However, it appears that not all casting swims are successful, as many spontaneous swims terminate after only a few seconds without leading to sustained taxis swims.

The necessity of directional shading of the photoreceptors is evident in larvae unable to synthesize pigment (melanin) due to a mutation in the tyrosine hydroxylase gene (*10, 11*). While these larvae initiate spontaneous swims and their photoreceptors remain functional, all spontaneous swims quickly terminate, and phototaxis is never observed. Similarly, when wild type larvae are observed in the absence of visual cues (*i*.*e*., under far-red illumination) phototaxis is absent, but short spontaneous swims persist. Spontaneous swims occur at a frequency of 2.6 + 1.1 swims/minute, while treating larvae with the GABAA receptor antagonist picrotoxin increases spontaneous swim frequency to 7.2 + 2.8 swims/minute, indicating that initiation of spontaneous swims is under inhibitory regulation (*6*). Spontaneous swims are even more profoundly disrupted in larvae homozygous from the forebrain mutant *frimousse* (*frm/frm*) [Figure 1A; (*12, 13*)]. Two distinct behavioral phenotypes are observed in *frm/frm* larvae. The majority display only short spontaneous swims, but with increased frequency (8.1 + 5.1 swims/minute), as well as with greatly reduced variability in the interval between swims (*6*). A subset of *frm/frm* larvae show long spontaneous taxis-like swims, but which occur in the absence of visible light. We report here that among the forebrain defects of *frm/frm* larvae is the absence of a single slow autonomously oscillating inhibitory neuron. In wild type larvae this inhibitory neuron targets the phototaxis circuit. Ablation of this neuron results in spontaneous long swims resembling taxis swims, but which occur independently of visual cues. The expression of the transcription factor *Otp* by this neuron suggests shared origins with oscillating hypothalamic neurons.

## RESULTS

### Inhibitory neurons in the forebrain

We previously reported that the *Ciona* larval midbrain contains a cluster of VGAT positive (VGAT^+^) neurons but we failed to identify VGAT^+^ neurons in the forebrain outside of a subset of photoreceptors (*6*). However, upon closer inspection, we found two additional VGAT^+^ forebrain neurons (provisionally named fb1 and fb2; Fig. 1B). These two neurons, as well as the VGAT^+^ photoreceptors, were absent in *frm/frm* larvae (Fig. 1C and (*13*)). fb1 and fb2 were also observed in a stable transgenic line expressing Kaede fluorescent protein under the control of the VGAT promoter [Fig. 1D; (*14*)]. The ventral neuron (fb2) is the same as the previously described Otp-expressing VGAT^+^ neuron (Fig. 1E; Fig. S1A) that was identified with a reporter plasmid containing the Otp *cis*-regulatory region (*15, 16*). We also observed expression of the Otp reporter in three of the seven *ascending motor ganglion* neurons in the hindbrain (AMGs; specifically, AMGs 2, 3 and 4; Fig. 1E). An *in situ* hybridization for Otp in *Ciona* larvae shows much wider expression than observed with the reporter construct, including additional neurons in the forebrain (Fig. S1B).

We previously reported that the number and position of neurons, as assessed by *in situ* hybridization, is consistent between *Ciona* larvae, being nearly identical in the hindbrain and highly similar in the midbrain (*6*). These observations allowed the mapping of 3-dimensional fluorescent *in situ* expression datasets to the single 3-dimensional connectome reconstruction in order to predict neurotransmitter use within the CNS. By similar reasoning, fb2 was consistently observed in *Ciona* larvae and in a consistent location, indicating that this neuron should be present in the 177 neurons identified in the *Ciona* connectome. To identify fb2 among the connectome neurons, we registered 3-dimensional VGAT *in situ* datasets to the connectome reconstruction using known VGAT^+^ neurons as anchors, as described previously (*6*). Based on the registration, fb2 mapped to *coronet-associated brain vesicle intrinsic neuron 78* [cor-assBVIN78; (*8, 17*)] (Fig. 2A). The reconstruction of cor-assBVIN78 from the connectome electron micrographs is strikingly similar to fb2 in terms of the shape of the soma, length of the axon, and the highly branched axonal terminus (Fig. 2B and C). Due to known minor variations in neuron location between larvae, we also investigated the neurons surrounding cor-assBVIN78 as candidates, none of which strongly resembled fb2 (Fig. 3A). Three of the surrounding neurons could easily be ruled out (coronet neurons 7, 10 and prRN100), and the remaining candidate neurons are all BVINs but had morphologies different from fb2. Based on similar anatomical location and cellular morphology we conclude that cor-assBVIN78 is the most likely neuron of the connectome dataset corresponding to fb2. Moreover, a comparison of the synaptic targets of cor-assBVIN78 to those of the surrounding BVINs, as given by the connectome, shows that they vary greatly (Fig. 3B), with cor-assBVIN78 standing apart in its targeting of five of the six *photoreceptor relay neurons* (prRNs). As will be presented below, the targeting of the prRNs is consistent with the activity of fb2. Because of the convergence of support for the identity of fb2, we will refer to it as cor-assBVIN78 for the remainder of this manuscript.

**Figure. 2.**
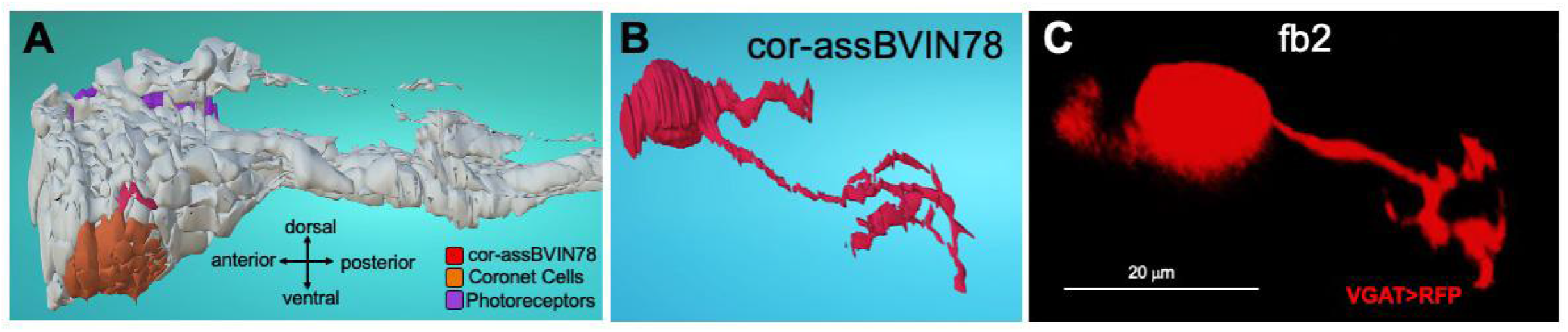
Mapping of fb2 to the *Ciona* connectome. **(A)** The location of neuron cor-assBVIN78 in the reconstructed CNS of the connectome larva. Also indicated are the coronet cells and the photoreceptors. **(B)** Reconstruction of neuron cor-assBVIN78 from the connectome. **(C)** Maximum intensity projection of confocal imaging of neuron fb2, labeled with RFP driven by the VGAT promoter. Images A and B are from the *Cionectome* connectome viewer (https://morphonet.org/2Tn9x6ZF).

**Figure. 3.**
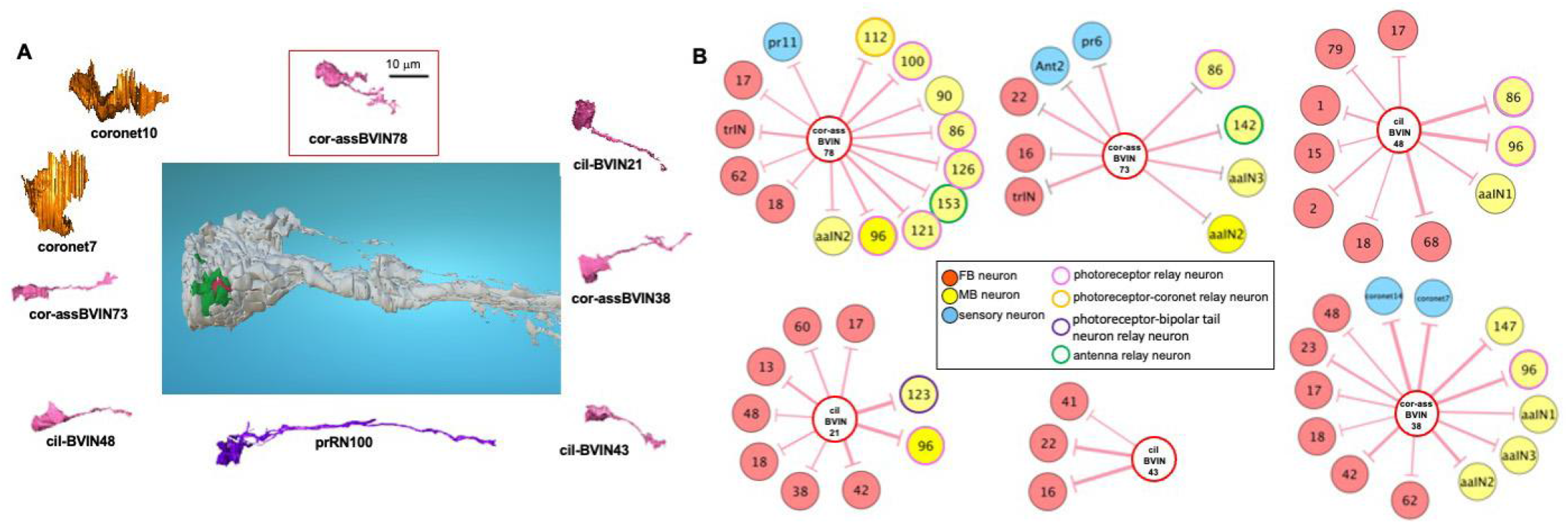
Candidate fb2 neurons. **(A)** The central panel shows the reconstructed *Ciona* CNS with cor-assBVIN78 in red and eight neighboring neurons in green. Reconstructions of each of the neurons from the connectome electron micrographs are shown. Images are from the Cionectome connectome viewer (https://morphonet.org/2Tn9x6ZF). **(B)** Chemical synaptic targets of the six candidate BVIN neurons. Target neurons are color coded depending whether they are sensory, forebrain (FB) or midbrain (MB), and numbered neuron IDs are according to (*17*). Relay neurons are circled and color coded according to their classification (*17*). (cil-BVIN, ciliated BV intrinsic interneurons; cor-assBVIN, coronet associated ciliated BV interneurons; prRN, photoreceptor relay neuron; aaIN, anaxonal arborizing neuron).

### cor-assBVIN78 is an autonomously oscillating neuron

We examined the calcium dynamics of cor-assBVIN78 as a proxy for neural activity using either VGAT or Otp *cis*-regulatory fragments to drive jGCaMP7f or jGCaMP8m (*18, 19*), and observed that cor-assBVIN78 consistently showed oscillating calcium transients (Fig. 4A). No other oscillating VGAT^+^ neurons were observed in larvae with GCaMP, although VGAT^+^ neurons with oscillation frequencies higher than can be resolved temporally with GCaMP remain a possibility. Examples of time-lapse recordings from three larvae are shown in Movie S2. While the cor-assBVIN78 consistently showed oscillatory behavior, we observed variation both in period and amplitude of GCaMP fluorescence peaks both within, and between, larvae (Figs. 4B and 5). Figure 4B shows plots of the 1/interpeak GCaMP intervals for 10 larvae, used as an estimate of variation in oscillation frequencies. The mean peak intervals calculated for nine of the larvae clustered in the range from 0.36 to 1.06 sec^-1^, with one outlier (larva I) having a mean at 2.58 sec^-1^. Pairwise comparisons showed statistically significant differences in peak intervals between larvae (Fig. 4C; Dunn’s test). Time and frequency domain graphs, as well as autocorrelation, of the cor-assBVIN78 GCaMP fluorescence data for the ten larvae are presented in Fig. S2, with the dominant oscillation frequencies indicated (except for larva E, whose dominant frequency could not be distinguished from the low frequency noise). The average oscillation frequency (excluding larvae E and I) was 0.61 ± 0.29 Hz (n = 8 larvae). The oscillation frequency of cor-assBVIN78 in larva I at 2.99 Hz was much higher than in all other larvae.

**Figure 4.**
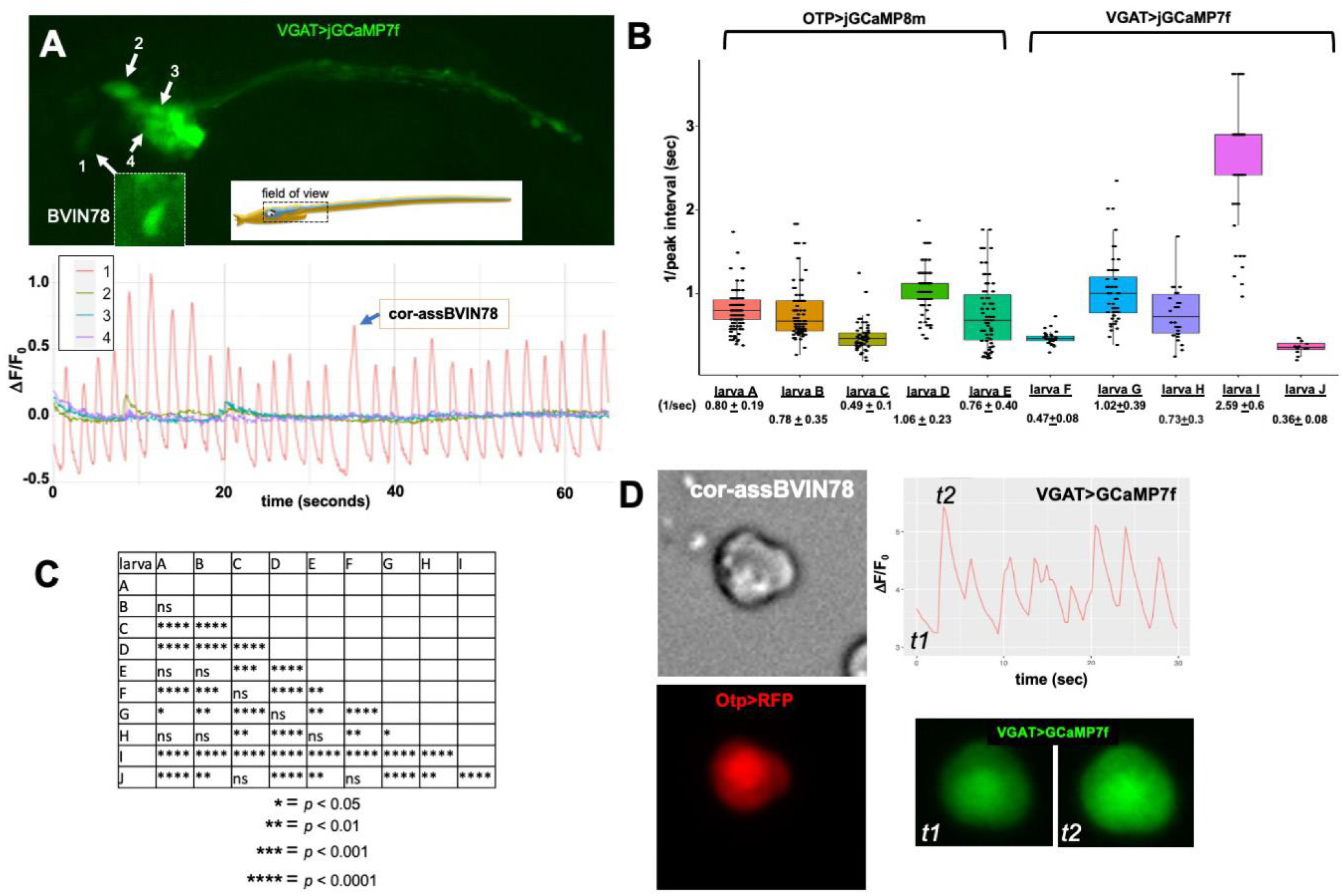
Activity of cor-assBVIN78. (**A**) Calcium imaging of cor-assBVIN78 using jGCaMP7f driven by the VGAT promoter (VGAT>jGCaMP7f). The chart shows relative changes in fluorescence over time for the four cells indicated, including cor-assBVIN78 (cell1). (**B)** Box plots of 1/interpeak intervals for five Otp>jGCaMP8m- and five VGAT>jGCaMP7f-expressing larvae (A-J). **(C)** Pairwise comparisons of the interpeak intervals for the ten larvae shown in panel B. p-values were adjusted by the Holm method. **(D)**. Isolated cor-assBVIN78 neuron expressing Otp>RFP and VGAT>jGCaMP7f. Chart shows normalized GCaMP7f fluorescence for the isolated cell. The center line is the median. The lower and upper box limit correspond to the first and third quartiles. The upper and lower whisker extends from the box limit to the largest value no further than 1.5 * IQR from the box limit. All data points were shown. See also Figure S2, Figure S3, Movie S2 and Movie S3.

We next examined whether cor-assBVIN78 retains oscillating activity when cultured in isolation. For this experiment we took advantage of the fact that the transient transfection method used here [electroporation at the one-cell stage (*20*)] can result in mosaic expression. This allowed us to manually select for larvae expressing the Otp reporter plasmid only in cor-assBVIN78 (*e*.*g*., larva shown in Fig. S1A). Larvae selected this way, and co-expressing Otp>RFP and VGAT>jGCaMP7f, were dissociated to single cells as described previously (*21*). The dissociated cell suspension was settled on a microscope slide under a coverslip, and cells co-expressing RFP and jGCaMP7f were identified. Calcium imaging showed that the isolated cor-assBVIN78 neurons continued to oscillate in isolation (Fig. 4D; and Movie S3). We collected jGCaMP7f data on four isolated cor-assBVIN78 neurons (Fig. S3). Fourier analysis for three of the cells each yielded a single dominant frequency that averaged 0.23 ± 0.06 Hz (n = 3 cells), while the data from the fourth cell did not resolve to a single dominant frequency. The slower oscillation of the isolated cor-assBVIN78 neurons compared to these neurons in intact larvae could be the result of their isolation from other modulating neurons (see Fig. 5, below). However, it is also possible the slower oscillation could be due, at least in part, to the medium used to observe the isolated cor-assBVIN78s (seawater) is not optimal.

**Figure. 5.**
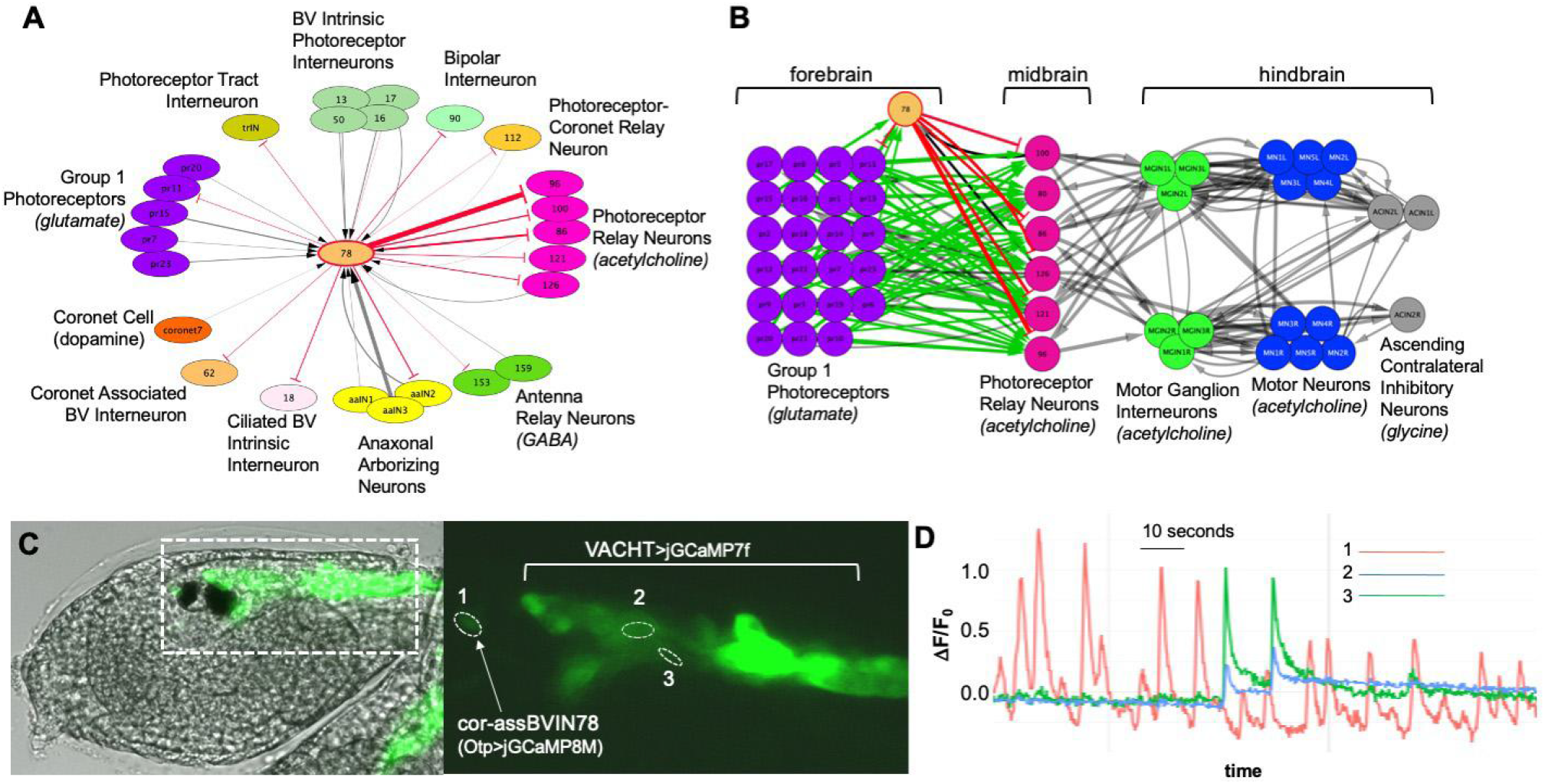
Connectivity and activity of cor-assBVIN78. **(A)** Synaptic connectivity of cor-assBVIN78 (shortened to *“78”* in panels A and B) as given by the *Ciona* connectome. Gray lines indicate synaptic inputs and red lines indicate synaptic outputs. **(B)** Minimal phototaxis circuit with the inhibitory inputs from cor-assBVIN78 shown in red. Green lines indicate projections from photoreceptors. All other synaptic connections are in gray. **(C)** Expression of jGCaMP8M from the Otp and VACHT regulatory elements in the larval CNS. **(D)** Normalized calcium fluorescence for the circled regions of interest (ROIs) in panel C. ROI1 corresponds to cor-assBVIN78. ROIs 2 and 3 appear to be the soma and axon of the same VACHT^+^ neuron, respectively. The thickness of the lines in panels **A** and **B** is proportional to the physical size of the synapse as determined from electron micrographs. See also Movie S4.

### Synaptic Connectivity and Activity of cor-assBVIN78

The complete chemical synaptic connectivity of cor-assBVIN78, as given by the connectome, is shown in Fig. 5A, with outputs from cor-assBVIN78 in red, and inputs to cor-assBVIN78 in gray. The connectome quantified the physical sizes of synapses as a putative measure of synaptic strength (*8*), and this is reflected in the thickness of the lines in Fig. 5A and B. Accordingly, the major synaptic targets of cor-assBVIN78 are the *photoreceptor relay neurons* (prRN) of the midbrain, which also receives input from the Group I photoreceptors (Fig. 5B). The relay neurons are so-named because they project from the midbrain to the hindbrain. The connectome identifies 38 neurons as being relay neurons, six of which are prRNs, with five of these receiving synaptic input from cor-assBVIN78. While cor-assBVIN78 is identified as being coronet cell-associated, it receives only weak input from a single coronet cell. However as a neuron class, the *corr-assBVIN* cells are characterized by their ciliary projections that contact the coronet cells (*17*). The synaptic inputs to cor-assBVIN78 also tie it to photoreception, as it receives input from four Group I photoreceptors (PRIs), as well as from brain intrinsic interneurons that receive photoreceptor input [neurons 13, 17, 50 and 16; (*8*)]. Fig. 5B places cor-assBVIN78 in the context of the minimal phototaxis circuit (*6, 8*), which starts with the Group I photoreceptors projecting to the midbrain prRNs, which in turn project to the left and right hindbrain *motor ganglion interneurons* (MGINs). The MGINs then project to motor neurons on their respective sides. Twenty-one of the twenty-three PRIs are glutamatergic, one is GABAergic, and one is dual GABAergic and glutamatergic. The significance of the single GABAergic and GABAergic/glutamatergic photoreceptors is not known, but they were found in all larvae examined (*6*). The prRNs are predicted to be primarily cholinergic, and thus this circuit appears to be an excitatory relay to the cholinergic MGINs (*6*). By contrast, cor-assBVIN78, due to its expression of VGAT, is predicted to be inhibitory, and its synaptic input to the prRNs suggest a gating function in the phototaxis pathway. Evidence for gating by cor-assBVIN78 was gained by simultaneous calcium imaging of cor-assBVIN78 and cholinergic midbrain neurons (Fig. 5C and D). During calcium imaging, we occasionally observe spontaneous calcium transients in VACHT^+^ midbrain neurons. As shown in Fig. 5, spikes in VACHT^+^ neurons correspond to troughs in cor-assBVIN78 calcium activity (see also Movie S4). We assume this is a prRN neuron, although we do not at present have a molecular marker that distinguishes the six prRNs from other midbrain relay neurons. However, our published quantification of *in situ* hybridization results from multiple larvae found only 11±1 VACHT^+^ midbrain neurons (*6*), which together with the connectivity data shown in Fig. 5A, is consistent with this spiking VACHT^+^ neuron being a prRN. We also observed examples of spontaneous spikes in midbrain VACHT^+^ neurons that do not correlate with cor-assBVIN78 activity (not shown).

### Ablation of cor-assBVIN78 results in long sustained spontaneous swims

Further evidence for gating activity of cor-assBVIN78 in the phototaxis pathway comes from loss-of-function analysis. cor-assBVIN78 was ablated in larvae carrying the stable VGAT>Kaede transgene using a UV laser. cor-assBVIN78 was identified based on Kaede fluorescence, then an area restricted to cor-assBVIN78 was selected and scanned with a UV laser at high intensity for ablation. Mock-ablated larvae were treated the same except a scanning area of the same size was chosen either anterior or posterior to cor-assBVIN78. Additional controls were treated as above, but received no ablation. As shown in Fig. 6A, cor-assBVIN78 was effectively ablated by this approach and only fluorescent debris was observed in the area of the ablation (circle in right panel). Following ablations, larvae were recovered from the slides, and swimming behavior was recorded in 3-minute sessions using only far-red illumination. The larvae from one experiment were combined in a single arena, and a previously described custom software script was used to track individual larva (*6*). Figure 6B and Movie S5 show combined swims for all larvae in the arena (7 control; 6 mock-ablated; and 9 cor-assBVIN78-ablated), with each swim a separate color, while Fig. S4A shows the swim tracks for individual larvae. A repeat experiment of the mock and cor-assBVIN78 ablations with a separate group of larvae is shown in Fig. S5 and Movie S6, with individual swims shown in Fig. S4B. For comparison of swimming behavior of ablated and mock-ablated larvae to normal larvae, we also collected a dataset of swims from unmanipulated larvae recorded both under far-red illumination and in the presence of directional visible light, which evokes phototaxis (*6, 10*). Representative swim traces from this group are shown in Fig. S6.

**Figure. 6.**
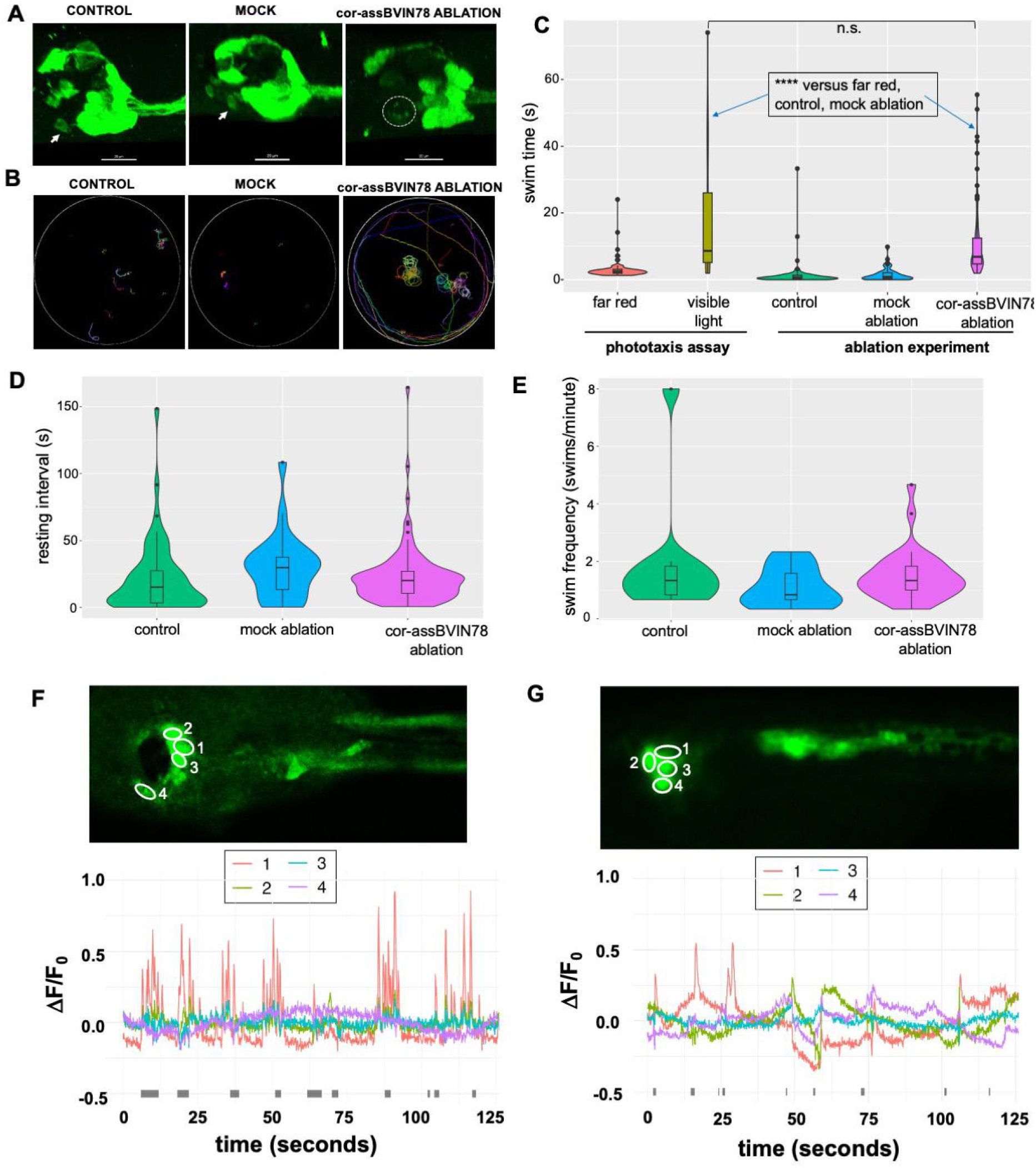
Ablation of cor-assBVIN78. **(A)** Laser ablation of cor-assBVIN78 in larvae stably expressing VGAT>Kaede. Arrows in “Control” and “Mock” indicate cor-assBVIN78. The circle in the right panel (cor-assBVIN78 ABLATION) indicates cell debris. Samples were immunostained with anti-Kaede antibody before confocal microscopy. **(B)** Swim tracks from eight CONTROL, six MOCK ablated, and nine cor-assBVIN78-ABLATED larvae. Each swim is assigned an arbitrary color. Swims for individual larvae are shown in Fig. S4. **(C)** Violin plots of the duration for swims from a phototaxis assay of unmanipulated larvae (left), and unablated (control), mock ablated, and cor-assBVIN78-ablated, larvae (right). (Dunn’s test; ****=*p*_*adj*_ <0.001; n.s., not significant). **D**. Violin plots of the resting interval between swims in control (unmanipulated larvae), mock ablated and cor-assBVIN78-ablated larvae. P>0.05 for all comparisons. **E**. Violin plots of swim frequency (swims/minute) in control (unmanipulated larvae), mock ablated and cor-assBVIN78-ablated larvae. P>0.05 for all comparisons. **F** and **G**. Plots of VACHT-jGCaMP7f in cor-assBVIN78-ablated, and mock-ablated, larvae, respectively. The numbered circled regions in the top panel correspond to the traces shown in the plots. Gray bars along the x-axis indicate periods in which tail beating was observed. See also Figure S4, Figure S5, Figure S6, Movie S5 and Movie S6.

The most obvious difference between the control groups (no ablation and mock ablation) and the cor-assBVIN78-ablated group, is the presence of long sustained swims in the latter group (Figure 6B and C; and Figs. S4 and S5). The long swims in the cor-assBVIN78-ablated larvae were observed despite the absence of visual cues *(i*.*e*., they were recorded under far-red light). Significantly, cor-assBVIN78 ablation did not result in continuous swimming. Rather, the larvae swam in spontaneous bouts that averaged 10.8 + 10.4 s (n= 98 swims). This was not significantly different from the duration of swims observed for normal larvae during phototaxis (18.5 + 19.5 s; n =24 swims; *p*_*adj*_ =0.52, Dunn’s test). The swims from both the cor-assBVIN78-ablated and normal phototactic swims were significantly longer (*p*_*adj*_<10^−6^, Dunn’s test) when compared to the three control groups: mock ablated, control (manipulated, but not ablated), and unmanipulated, which averaged 1.9 + 2.1 s (n= 33 swims), 1.6 + 4.4 s (n= 62 swims) and 3.2 + 3.3 s (n=68 swims), respectively (all controls recorded under far-red light). Thus ablation of cor-assBVIN78 evokes swims of similar length to those observed in phototaxis. Moreover, in both the cor-assBVIN78-ablated and the phototaxis swims we observed a mixture of both straight and tortuous swims (Fig. 6B and Fig. S6). We also analyzed the ablation dataset for the length of the resting intervals between swims, and the frequency of swims (Figure 6 D and E). We found no significant differences in these parameters between the cor-assBVIN78 ablated, mock ablated, and unablated control larvae. The results also indicate that cor-assBVIN78 ablation phenocopies one aspect of the behavioral phenotype of *frm/frm* larvae. In *frm/frm* we observed that the majority of larvae exhibited only short spontaneous swims - but with a higher frequency and lower variability. However, and similar to cor-assBVIN78-ablated larvae, a subset of *frm/frm* larvae exhibited long swims in the absence of visual cues (in fact, *frm/frm* larvae have no photoreceptors). The *frm* mutation is much more extensive than the simple loss of cor-assBVIN78, and results in the transfating of the entire forebrain to epidermis (*12, 13*). Nevertheless, the behavior of long-swimming *frm/frm* larvae is consistent with the removal of an inhibitory input from the forebrain. The reason why only some *frm/frm* larvae are able to initiate long swims suggests variable penetrance of the *frm* phenotype, with differing degrees of CNS disruption. Whether the absence of cor-assBVIN78 also plays a role in the altered frequency of short spontaneous swims in *frm/frm* larvae is not readily discernible from the results of the ablation. The rhythm of spontaneous swims is best observed in wild-type larvae recorded under far red illumination, or in *frm/frm* larvae. In both cases, long taxis-like swims are rare, and the short spontaneous swims lasting 1-2 seconds, and occurring every 9-15 seconds on average, are easily quantified. In cor-assBVIN78-ablated larvae we observed almost exclusively long swims lasting tens of seconds, so that the underlying short spontaneous swim frequency could not be quantified.

Labeling of cor-assBVIN78 with transgenic reporters shows that it projects to the midbrain (*e*.*g*., Figure 1E), and the connectome predicts that primary targets in the midbrain are the prRNs (Figure 5B). Thus, we predict that the behavioral phenotype resulting from cor-assBVIN78 ablation will be accompanied by increased firing of the prRNs. While we do not have a driver that is specific for the prRNs, they are predicted to make up approximately half of the ∼11 midbrain VACHT^+^ neurons (*6*). To determine if elevated firing of VACHT^+^ midbrain neurons in cor-assBVIN78-albated compared to mock-ablation larvae could be detected, embryos were first electroporated with Otp>RFP and VACHT>jGCaMP7f. At the larval stage, cor-assBVIN78 was ablated, as above, but using the Otp>RFP fluorescence as a guide. Figures 6F and G show plots of GCaMP fluorescence captured in 2-minute imaging sessions for cor-assBVIN78-albated, and mock-ablation larvae, respectively. Because the VACHT^+^ neurons are clustered in the midbrain, we could not resolve individual neurons, rather we analyzed the indicated region of interest (ROIs 1-4 in each panel) that cover the VACHT^+^ neurons in the midbrain. Consistent with our predictions, extensive spiking was observed in the cor-assBVIN78-albated larva (particularly ROI 1; panel F). The extent of spiking in the VACHT^+^ neurons in the mock-ablated was much less extensive (panel G).

## DISCUSSION

### Identity of fb2

The cross-identification of a single neuron between disparate datasets, such as between a single serial-section EM reconstruction and light microscopy images, as presented here, is inherently problematic. While light microscopy images of multiple larvae showed a consistent location for *fb-2* in the CNS, the fact we have only one connectome dataset limits the confidence of our 3-dimensional registration approach. Likewise, our results highlighting the similar cellular morphology of *fb-2* to cor-assBVIN78, but not to other nearby neurons, suffer from this same limitation. The ablation results, which are in agreement with predictions from the single connectome, would also be potentially strengthened by additional connectomic data. Nevertheless, and while acknowledging these limitations, we feel that the totality of the data make cor-assBVIN78 the only reasonable candidate for *fb-2*.

### *BVIN 78 and the* Ciona *forebrain*

The parallel anatomy, gene expression, and function of tunicate larval and vertebrate nervous systems has been extensively documented (*2*). The most anterior part of the *Ciona* CNS shows hallmarks that link it to a hypothetical proto-forebrain structure in the common ancestor of vertebrates and tunicates. These hallmarks include, for example, the expression of the transcription factors *Otx* and *Dmrt* (*4, 22, 23*). *In situ* hybridization surveys for markers of small-molecule neurotransmitter use show little or, for some markers, no expression in larval forebrain neurons. The few expressing neurons are the photoreceptors, which express both VGLUT and VGAT (*6, 24*), the coronet cells which appear to be dopaminergic (*25*) and, as reported here, *fb1* and *fb2* which express VGAT. For the remaining forebrain neurons, one possibility is that many are peptidergic (*26*) or, alternatively, that they are quiescent with respect to neurotransmitter use until after metamorphosis. Nevertheless, the connectome predicts extensive synaptic connectivity between *Ciona* forebrain neurons, as well as extensive descending projections (*8*). Despite this, and in contrast to the *Ciona* larval midbrain, which has a clear role as an integrating and relay center for sensory inputs (*3, 6, 7*), the function of *Ciona* forebrain remains poorly characterized. Among the *Ciona* larval forebrain neurons with known function are the photoreceptors, but they, in fact, originate more posteriorly, and then migrate to the forebrain (*27*). The coronet cells have attributes of sensory cells. While their function is unknown, they have been speculated to play a role in circadian rhythms (*16*). Thus, cor-assBVIN78 is the first non-sensory neuron of the *Ciona* forebrain with a known function.

### cor-assBVIN78, spontaneity, and phototaxis

In *Ciona* larvae, negative phototaxis initiates with spontaneous casting, which can then transition to a phototaxis swim - if a directional light cue is received (*10*). The results presented here indicate that cor-assBVIN78 acts on this transition. A plausible neural circuit for phototaxis in *Ciona* has been proposed, and consists of a relay from glutamatergic photoreceptors to motor neurons via two classes of cholinergic interneurons [Fig. 5B; (*6, 8*)]. The presence of an inhibitory neuron, particularly a slow-oscillating one, acting on the prRNs was not predicted by the circuit model. Nevertheless, loss of this neuron has profound behavioral effects. The results presented here provide several clues to the role of cor-assBVIN78 in the phototaxis circuit. While our results show that the ablation of cor-assBVIN78 is sufficient to evoke sustained taxis-like swims, we do not observe spontaneous swims in intact larvae corresponding to the rhythm of cor-assBVIN78. In other words, the troughs in cor-assBVIN78 inhibition, which occur at an average frequency of 0.62 Hz, alone are not sufficient to evoke swimming. This suggests that the inhibitory action of cor-assBVIN78 may act as a filter to suppress spurious inputs to the phototaxis circuit. However, this does not explain the oscillatory activity of cor-assBVIN78. Data presented here indicate that the oscillating inhibition by cor-assBVIN78 gates the activity of the photoreceptor relay neurons, making them preferentially responsive to phototactic cues arriving during troughs (Fig. 7). Observations of spontaneously firing cholinergic relay neurons support this hypothesis (Fig. 5C and D). The net effect of cor-assBVIN78 activity appears to add an inhibitory element to the transition from casting to taxis. This may explain why many casting swims do not transition to phototactic swims (*10*). Moreover, the rhythm of the spontaneous swim generator at ∼2.6/minute (0.43 Hz) is not synchronized with cor-assBVIN78 oscillation and, moreover, the two appear to act in opposition. Thus, their interaction should add an element of noise, or uncertainty, to the frequency of phototaxis, which would otherwise be a more predictable rhythmic behavior driven by the spontaneous swim generator. Thus, the oscillatory action of cor-assBVIN78 may impart a selective advantage to larvae by modifying phototaxis so as to make it less overtly rhythmic, and thereby less predictable, possibly helping to avoid predation. The variation observed in the frequency and amplitude of the oscillations, both within and between larvae, may accentuate this effect.

**Figure 7.**
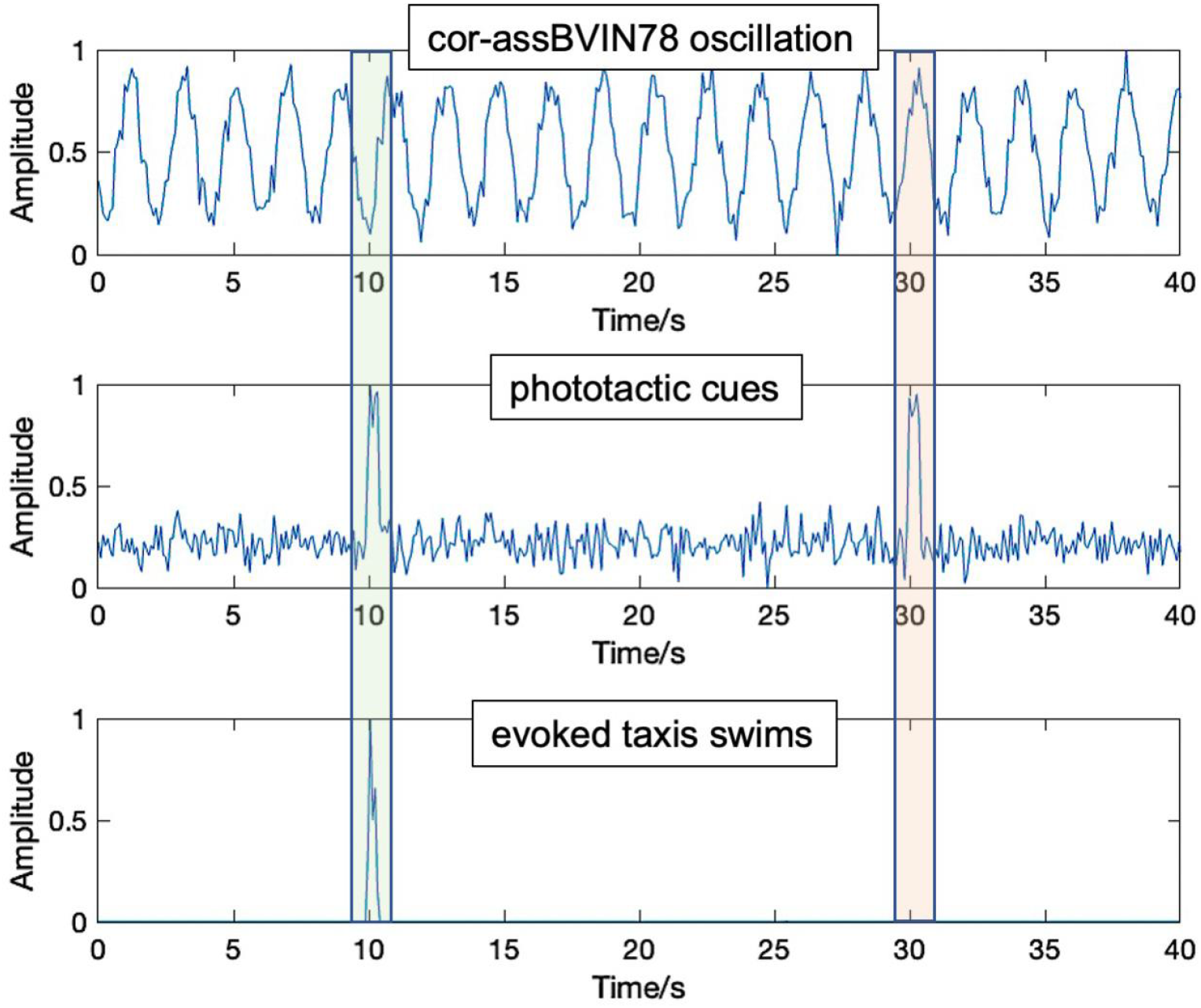
Model for cor-assBVIN78 gating of phototaxis. Hypothetical data illustrating the proposed filtering and gating activity of cor-assBVIN78 in the phototaxis circuit. Phototaxis cues generated via casting swims that correspond temporally to troughs in cor-assBVIN78 oscillations can evoke taxis swims (green box), while those corresponding temporally to peaks do not (red box). In addition, small fluctuations, indicated by the added noise to the phototactic cue graph, may not evoke swims even when corresponding temporally to troughs due to the proposed filtering activity of cor-assBVIN78.

### Relationship of cor-assBVIN78 to the central pattern generator

*Ciona* larvae alternate activation of the left and right flanks of tail muscles to generate rhythmic swimming. Components of the central pattern generator (CPG) have been identified in *Ciona* larvae, and include left and right pairs of contralaterally projecting inhibitory (glycinergic) neurons that target the right and left *motor ganglion interneurons* (MGINs) of the hindbrain, respectively (*28*). In addition, one left/right pair of the ten hindbrain motor neurons has been shown to have autonomous oscillating behavior that determines swim rhythmicity (*5, 29*). Moreover, recent experiments in which larvae were cut into progressively smaller segments demonstrated that a fragment containing only the hindbrain and an anterior segment of the spinal cord (referred to as the caudal nerve cord) was able to autonomously initiate periodic and rhythmic swim bouts (*30*). Taken together, these results point to the central pattern generator as residing exclusively in the hindbrain and spinal cord. The physical location and slow oscillation of cor-assBVIN78 suggest it is not a component of the CPG, but rather acts upstream. In addition, we observed that ablation of cor-assBVIN78 did not result in continuous swimming. Rather, the ablated larvae swim in discrete bouts, similar to phototactic swims. Thus, while cor-assBVIN78 appears to act to temporally inhibit the initiation of swim bouts, it does not appear to be essential for their termination. Perhaps the mechanisms determining the persistence and termination of evoked swim bouts may be inherent to the CPG.

### cor-assBVIN78 and other chordates

With fewer than 50 neurons, the diminutive size of the *Ciona* larval forebrain relative to those of vertebrates presents an opportunity to explore the function of individual neurons, as is done here for cor-assBVIN78. However, linking individual cell types between clades as distantly related as tunicates and vertebrates is more challenging. Nevertheless, the anatomical location, expression of *Otp*, oscillatory activity, and synaptic connectivity with the visual system all point to a link between cor-assBVIN78 with an ancient chordate neuronal type that also went on to populate the vertebrate hypothalamus (*31, 32*). The neighboring dopaminergic coronet cells of the *Ciona* forebrain are also thought to have ties to the hypothalamus (*16*). Oscillatory behavior in the hypothalamus has been studied predominantly in the context of regulating circadian cycles (*31*). However, it is unlikely that cor-assBVIN78 is involved in circadian rhythms in *Ciona* larvae. The larval stage is brief in *Ciona*, lasting only one or two days, following which the animal undergoes metamorphosis, which includes an extensive remodeling of the nervous system (*33*). Moreover, no circadian behavior has been described in *Ciona* larvae, and the *Ciona* genome appears to have lost many of the canonical clock genes, including *clock* and *per* (*34, 35*). Rather, as we proposed above, cor-assBVIN78 appears to have a role in regulating rhythmic taxis behavior. Whether this is a unique mechanism to *Ciona* (or tunicates), or is more widespread, remains to be determined.

## Data Availability

All data from this manuscript is deposited in the Brain Image Library (https://www.brainimagelibrary.org/index.html). All code from this manuscript is hosted at GitHub (https://github.com/CionaLab/phnn).

## Acknowledgements

We thank Yasunori Sasakura for transgenic *Ciona* lines, and Robert Zeller for plasmids. We also thank Kerrianne Ryan for helpful discussions. Emmanuel Faure thanks Patrick Lemaire for financial support. We also acknowledge the support provided by the ANISEED web portal. We acknowledge our funding sources: DE-SC0021978 from Department of Energy Office of Science to WCS; R34 NS127106 from NIH/NINDS to WCS; UCSB CCS Create Fund Summer Fellowship to JM; and ANR-21-NEUC-0004 to Patrick Lemaire.

## Author contributions

Conceptualization: J.C, E.N-S, M.K and W.S

Methodology: J.C, E.N-S, M.K and W.S

Investigation: J.C, E.N-S, M.K, J.M and W.S

Formal Analysis: Y.M

Data Curation: Y.M.

Writing--Original Draft: W.S.

Writing--Review and Editing: J.C, E.N-S, M.K, Y.M, C.B, W.S, T.L, B.G and E.F.

Funding Acquisition: W.C.

Software: C.B, T.L, B.G and E.F.

## Star methods

### Resource availability Lead contact

Further information and requests for resources and reagents should be directed to the lead contact, William C. Smith.

### Materials availability

Plasmids generated in this study are available upon request.

### Data and Code availability

All images and time lapse data generated for this paper have been deposited at the Brain Image Library (https://brainimagelibrary.org/). DOIs will be listed in the key resources table when they are available. All original code has been deposited at Github and Zenodo and is publicly available as of the date of publication. DOIs are listed in the key resources table.

Any additional information about the data reported in this paper is available from the lead contact upon request.

### Experimental model and subject details

Adult *Ciona robusta* were collected at the Santa Barbara Harbor or cultured in the UC Santa Barbara Marine Laboratory, a continuous-flow seawater system. Larvae were generated by dissecting gametes from adults. VGAT>Kaede transgenic larvae were produced by crossing wild type eggs with frozen VGAT>Kaede sperm aliquots. Electroporated embryos were first dechorionated before being electroporated with the construct of interest. All embryos were incubated at 18°C for optimal growth conditions.

### Key Resources Table

| REAGENT or RESOURCE | SOURCE | IDENTIFIER |
| --- | --- | --- |
| **Chemicals, Peptides, and Recombinant Proteins** |  |  |
| Sodium thioglycolate | Sigma | T0632 |
| Protease E | Sigma | P5147 |
| Trypsin | Sigma | T0303 |
| collagenase | Sigma | C9891 |
| MS222 | Sigma | E10521 |
| Ultra low melt agarose | Sigma | A5030 |
| **Antibodies and Probes** |  |  |
| VGAT *in situ* probe | Molecular Instruments | https://www.molecularinstruments.com |
| VACHT *in situ* probe | Molecular Instruments | https://www.molecularinstruments.com |
| OTP *in situ* probe | Molecular Instruments | https://www.molecularinstruments.com |
| Kaede anti rabbit | MBL | PM012M |
| goat anti-rabbit AlexaFluor488 | Invitrogen | A11034 |
| rabbit anti-GFP | Invitrogen | A11122 |
| **Experimental Models: Organisms/Strains** |  |  |
| *Ciona robusta* wild caught (formerly *Ciona intestinalis* type A) | Santa Barbara Harbor | N/A |
| *Ciona robusta* VGAT>Kaede transgenic | Ciona Intestinalis Transgenic Line Resources (CITRES) | Tg[MiCiVGATK]2 |
| **Recombinant DNA** |  |  |
| Otp>CiRFP | This manuscript | N/A |
| Otp>jGCaMP8m | This manuscript | N/A |
| VGAT>jGCaMP7f | This manuscript | N/A |
| VACHT>jGCaMP7f | This manuscript | N/A |
| **Software and Algorithms** |  |  |
| Original codes | https://github.com/CionaLab/phnn | 10.5281/zenodo.7996629 |
| Fiji | https://imagej.net/Fiji | 2.13.1 |
| MATLAB | https://www.mathworks.com/products/matlab.html | 2023a |
| R | https://www.r-project.org/ | 4.3.0 |
| Tidyverse | https://www.tidyverse.org | 10.21105/joss.01686 |
| rstatix | https://CRAN.R-project.org/package=rstatix | 0.7.2 |
| ggpmisc | https://CRAN.R-project.org/package=ggpmisc | 0.5.2 |
| **Other** |  |  |
| ORCA-Flash V4.0 C11440-22CU camera | Hamamatsu | C11440-22CU |
| Zoom 7000 lens | Navitar | ZOOM 7000 |
| 52mm red 25 filter | Tiffen | 52R25 |
| BioLED optical head, Precision LED Spot light, 700 nm | Mightex Systems | BLS-PLS-0700-01-S |
| BioLED optical head, Precision LED Spot light, 505 nm | Mightex Systems | BLS-PLS-0505—030-04 |
| BioLED light source control module | Mightex Systems | BLS-SA04-US |

## MATERIAL AND METHODS

### Animals and histological methods

Adult *Ciona robusta* (aka, *C. intestinalis* type A) were collected at the Santa Barbara Harbor with the exception of the stable VGAT>Kaede line (Tg[MiCiVGATK]2; obtained from CITRES https://marinebio.nbrp.jp/ciona/), which was cultured at the UC Santa Barbara Marine Laboratory. Hybridization chain reaction *in situ* was performed as described previously, including probes for VGAT and VACHT (*6*). The mRNA sequence to which the probe set was chosen for Otp included 1898 bases, beginning UUUUUAAGU (at 5’ end) and ending with UAAAUCCCC (3’). The immunostaining procedure was performed as described previously (*36*). For labeling of larvae expressing Kaede, an anti-Kaede (rabbit) primary (MBL) was used, followed by a goat anti-rabbit AlexaFluor488 (Invitrogen). Animals expressing GCaMP were labeled using a rabbit anti-GFP primary (Invitrogen) and anti-rabbit AlexaFluor488 secondary (Invitrogen).

### Dechorionation and Electroporation

Fertilized eggs were electroporated with expression plasmids as described previously (*20*), with the exception that eggs were dechorionated using either the sodium thioglycolate/protease method, or with 0.1% Trypsin (Sigma T4799) in 10 mM TAPS pH 8.2 in filtered seawater, as described in (*37*).

### Reporter Plasmids

To generate the Otp>CiRFP, the Otp *cis*-regulatory fragment, as described in (*16*), was amplified by PCR and ligated into Xho1 and Sph1 restriction sites of pSP72-1.27 mRFPci, which has a codon optimized RFP for *Ciona* expression (*38*). The published Otp *cis*-regulatory fragment includes the first 112 amino acids of Otp. This expression strategy did not result in a functional GCaMP, perhaps due to the extra amino acids. Therefore, to make Otp>jGCaMP8m, the native ATG of Otp was mutated using a gBlock (IDT) before jGCaMP8m was added. jGCaMP8m was amplified using the following primers which add a *Ciona*-specific Kozak sequence: CCtctagaCAGAAAAAATGacgcgtcgcaagaagaccttc and ttGAATTCttacttcgctgtcatcatttgtac. The plasmid jGCaMP8m was obtained from Addgene (#162381). VGAT>GCaMP7f was synthesized by Twist bioscience (https://www.twistbioscience.com/) and contains a 2.5Kb VGAT *cis*-regulatory fragment and jGCaMP7f with the above *Ciona* Kozak sequence. To generate VACHT>GCaMP7f, GCaMP7f was PCR amplified and inserted into pSPCiVACHTK (*14*).

### GCaMP imaging and statistical analysis

Larva expressing transgenes for VGAT>jGCaMP7f, Otp>jGCaMP8m or VACHT>jGCaMP7f were mounted under coverslips raised with ∼80 *μ*m spacers made with adhesive reinforcement labels (Avery, #5729) on glass slides in filtered seawater at 24-26 hour post fertilization. Larva were held stationary by removing excess water under coverslips. Time series of 1-3 minutes were acquired at a frame rate of 10-15 frames per second with a Leica DM6B fluorescence microscope with a 20X lens at 18C. Single cells were imaged at a frame rate of 3-5 frames per second for 30 seconds.

GCaMP movies were analyzed in ImageJ (*39*). After selecting regions of interest (ROIs), fluorescence intensity was measured. The baseline value of fluorescence (F0) was defined as 20% of the mean fluorescence for each ROI. F0 was then used to calculate ΔF/F0 as (F-F0/F0) (*18*). The artifactual drift in ΔF/F0 over time was removed by taking the residual from a linear model with ΔF/F0 as the dependent variable and time as the independent variable in R. Autocorrelation and Fourier transform were performed in MATLAB R2023a (https://www.mathworks.com) on the residuals to determine the dominant frequencies (*40*). The peaks were determined by finding the local maxima of ΔF/F0 using ggpmisc 0.5.2 (https://github.com/aphalo/ggpmisc). Kruskal-Wallis test was performed to detect the difference between the 1/peak intervals among the groups. One-tailed Dunn’s test was performed *post hoc* to determine the pairwise difference between groups. Holm–Bonferroni method was used for multiple comparison correction. Both tests were performed using rstatix 0.7.2 (https://rpkgs.datanovia.com/rstatix/). The groups contain the following number of peaks: A 142, B 87, C 57, D 124, E 71, F 29, G 56, H 30, I 63 and J 12. Data processing was performed with Tidyverse (*41*)in R 4.3.0 (https://www.R-project.org/), if not otherwise specified. For box plots, the center line is the median. The lower and upper box limit correspond to the first and third quartiles. The upper and lower whisker extends from the box limit to the largest value no further than 1.5 * IQR from the box limit. All data points are shown with horizontal jitters.

### Larva dissociation and cell imaging

Eggs were co-electroporated with Otp>RFP and VGAT>GCaMP7f expression plasmid. The resulting larvae were screened for those expressing RFP only in BVIN78 (and not in the AMGs) using a fluorescence stereomicroscope. At 26 hours post fertilization, the selected larvae were dissociated to single cells using a modified zebrafish cell dissociation protocol (*42*). Briefly, the larvae were placed in 500 μL dissociation mix (0.25% trypsin and 4 mg/ml collagenase calcium-free artificial seawater). Larvae were pipetted every 30 seconds at 30ºC until intact larvae were no longer visible (∼5 minutes). To stop the dissociation, 800 μL of calcium-free seawater was added and the sample was centrifuged for 5 minutes at 700 x g. The cells were washed twice with 1mL of calcium-free seawater, then resuspended in 20-40 μL of seawater and settled on a microscope slide under a coverslip for imaging on a LEICA DM6B fluorescence microscope.

### Laser Ablation of BVIN78

Wild type eggs were fertilized with sperm from a stable VGAT>Kaede transgenic transgenic line (*14*). Following hatching at ∼21 hour post-fertilization at 18C, the larvae were selected for VGAT>Kaede expression using a fluorescence stereomicroscope, anesthetized with 100μg/mL MS222, and then mounted individually on glass slides under coverslips raised with ∼50 *μ*m spacers strips cut from Nunc aluminum seal tape (ThermoFisher, cat. no. 232698). Using the epifluorescence function of an Olympus Fluoview1000 scanning confocal microscope, Kaede was first photoconverted to red using the UV filter cube. Then, with confocal scanning using the 546 laser and an optical zoom of 10-12x under the 40x oil lens, the cell to be ablated was first identified and a region of interest demarcated. Neuron BVIN78, or an adjacent cell in the forebrain (mock ablation), was selected. The selected cell was then ablated using a 405 nm laser at 95% intensity for 150 seconds. Unablated (control) larvae were treated with MS222 and mounted on slides. Larvae were then recovered from the microscopic slides, placed into a petri dish of seawater to recover from the MS222 for at least 30 minutes, and then assessed for spontaneous swims.

### Behavioral Assays

For imaging spontaneous swimming behavior, larvae at 25 hours post-fertilization were placed in an imaging arena consisting of 45 mm X 14 mm circular depression made in 60 mm petri dish of 1% agarose in seawater (*i*.*e*., the walls and floor were made of 1% agarose to prevent the larvae from sticking). The depression was filled with seawater containing 0.1% ultra low-melt agarose, which remains liquid but helps prevent the larvae from becoming caught to the air-water interface. The petri dish was placed on a stage and recorded under far-red (700 nm) LED illumination with Hamamatsu ORCA-Flash V4.0 C11440-22CU camera fitted with a Navitar Zoom 7000 lens. Swims were recorded in three-minute capture sessions at 10 frames per second. For imaging phototactic swims, larvae were recorded with a directional 505 nm light, as described previously (*6, 10*). Image capture was the same as above, but a larger arena was used for this assay (85 X 10 mm).

### Quantification and statistical analysis

Larval swims were analyzed from two different clutches of larvae at 25 hours post fertilization, and three minute recordings of spontaneous swimming were made for each clutch with two to three replicates for each group. Spontaneous swimming was assessed by identifying the starting and stopping frames for each swim bout. This assessment was done using a previously described automated tracker (*6*)with added manual annotations. The projected paths of each larva were visualized using the MTrackJ plugin to ImageJ (*43*). Kruskal-Wallis test and Dunn’s test were performed to determine the pairwise difference between groups as described in the previous section. For box plots, the center line is the median. The lower and upper box limit correspond to the first and third quartiles. The upper and lower whisker extends from the box limit to the largest value no further than 1.5 * IQR from the box limit. Outliers are shown as points.

**Movie S1**. Representative phototactic swim (magenta). Other short (non-phototactic) swims are also tracked. The behavior arena was illuminated directionally from the right.

**Movie S2**. Three *Ciona* larvae expressing jGCaMP7f under control of the VGAT promoter. The oscillating cor-assBVIN78s are indicated by arrows. The movie plays at 10X normal speed.

**Movie S3**. Isolated cor-assBVIN78 neuron expressing jGCaMP7f under control of the VGAT promoter. The movie plays at 10X normal speed.

**Movie S4**. Larvae co-expressing Otp>jGCAMP8m in cor-assBVIN78 (isolated cell at left) and VACHT>jGCaMP7f (the remainder of the fluorescence). A calcium transient in a VACHT>jGCaMP7f neuron is observed at ∼6 seconds in this 19-second movie. The movie plays at 10X normal speed.

**Movie S5**. Swimming tracks of the first group of control, mock-ablated and BVI 78-albated larvae over a three-minute recording session. Each swim is assigned an arbitrary color. The movie plays at 10X normal speed.

**Movie S6**. Swimming tracks of the second group of mock-ablated and cor-assBVIN78-albated larvae over a three-minute recording session. Each swim is assigned an arbitrary color. The movie plays at 10X normal speed.

