## Supplemental Data for "A single oscillating proto-hypothalamic neuron gates taxis behavior in the primitive chordate *Ciona*"

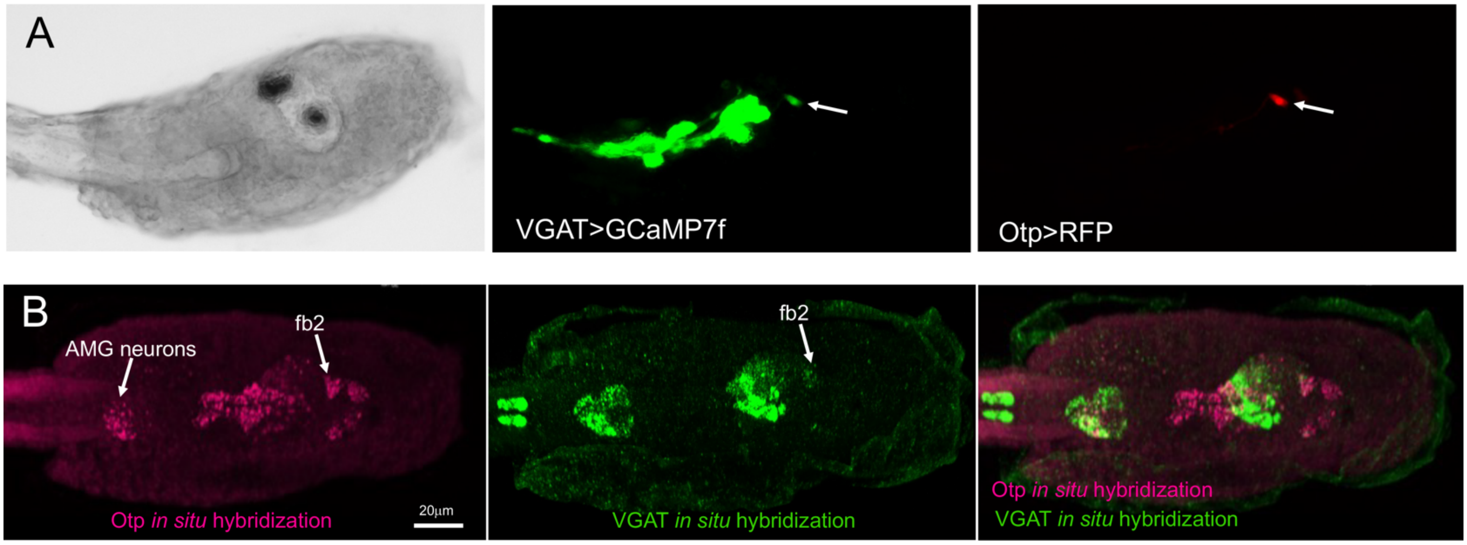


**Figure S1. Expression of Otp. Related to Figure 1. (A)** *Ciona* larva co-expressing VGAT>GCaMP7f and Otp>RPF to show overlap in fb2 (arrow). **(B)** Hybridization chain reaction (HCR) in situ hybridization in a *Ciona* larva for Otp (magenta) and VGAT (green). Image is a dorsal view of the trunk with anterior to the right.

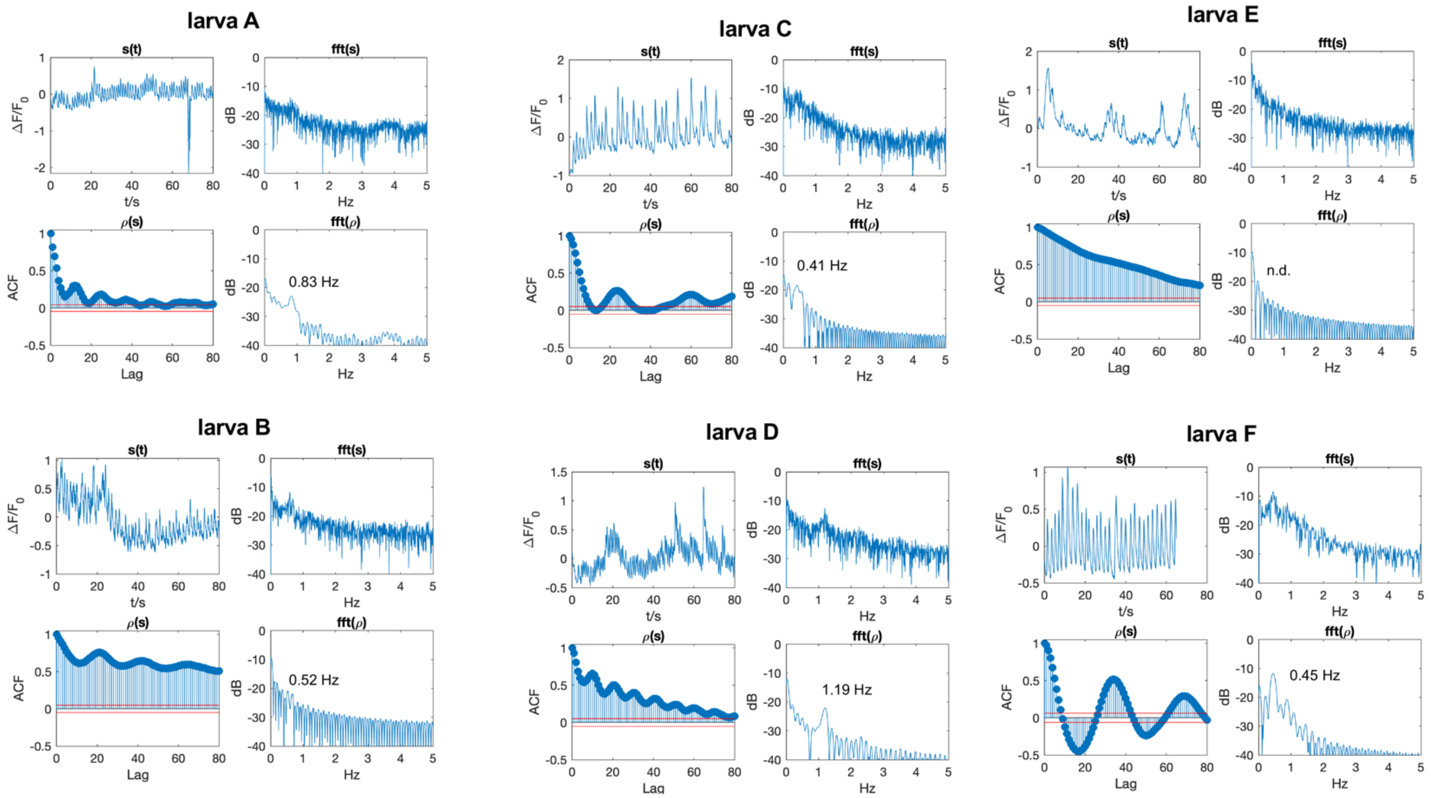


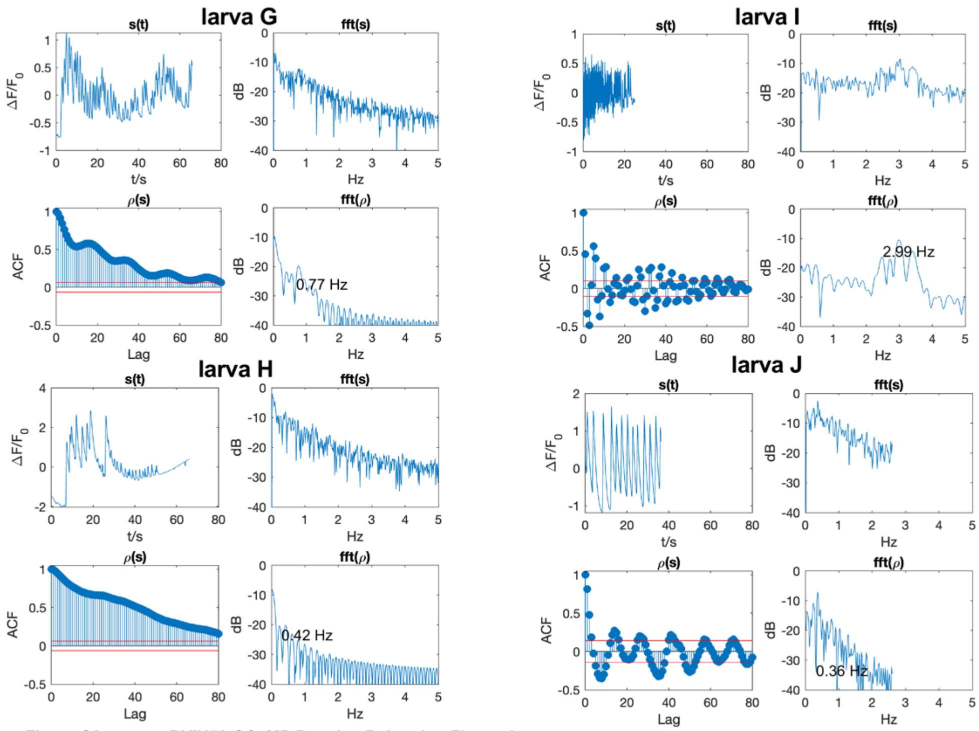


**Figure S2. cor-assBVIN78 GCaMP Results. Related to Figure 4.**
For each larva (A-J), the top left graphs show normalized GCAMP fluorescence ($\Delta F/F_0$) in cor-assBVIN78 for ten larvae. Larvae A-F were transfected with Otp>jGCaMP8m, and F-J were transfected with VGAT>jGCaMP7f. The top right graphs show the power spectra of the fluorescence signal after Fast Fourier Transform (FFT). The bottom left graphs show the Autocorrelation Function (ACF) until lag 80. The 95% confidence intervals are indicated by the two red lines. The bottom right graphs show the power spectra of the ACF. The dominant resonance frequency is indicated (except for larva E, which did not have a dominant frequency).

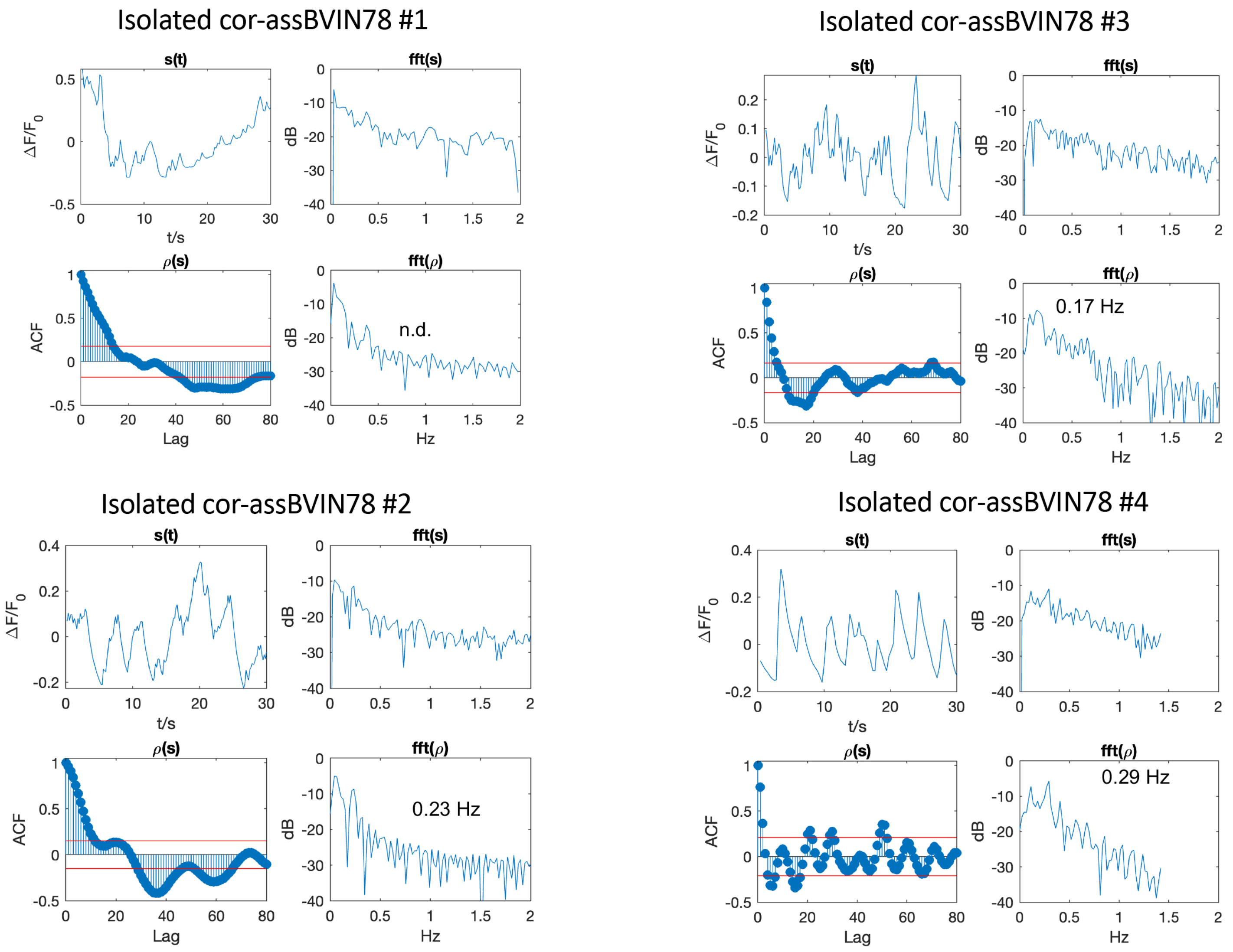


**Figure S3. GCaMP fluorescent in four isolated cor-assBVIN78 neurons. Related to Figure 4.** For each neuron, the top left graphs show normalized GCAMP fluorescence ($\Delta F/F_0$) from VGAT>jGCaMP7f. The top right graphs show the power spectra of the fluorescence signal after Fast Fourier Transform (FFT). The bottom left graphs show the Autocorrelation Function (ACF) until lag 80. The 95% confidence intervals are indicated by the two red lines. The bottom right graphs show the power spectra of the ACF. The dominant resonance frequency is indicated (except for cell 1, which did not have a dominant frequency).

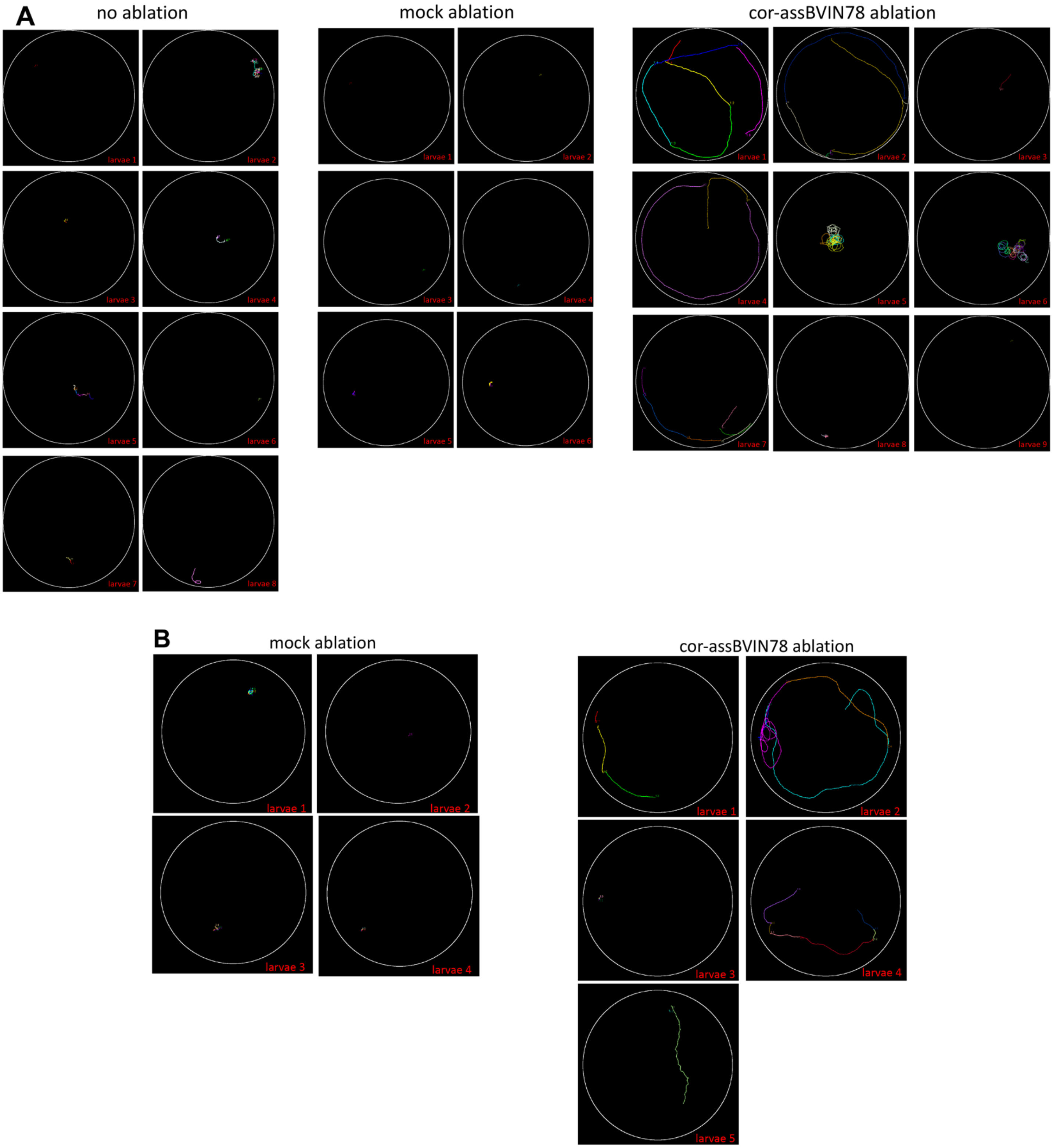


**Figure S4. Swim tracks of individual larvae. (A) Trial 1. (B) Trial 2. Related to Figure 6.** The swim tracks for 3-minute recording sessions are shown. Each swim is assigned an arbitrary color. Larvae received either no ablations, a mock ablation, or cor-assBVIN78 ablation.

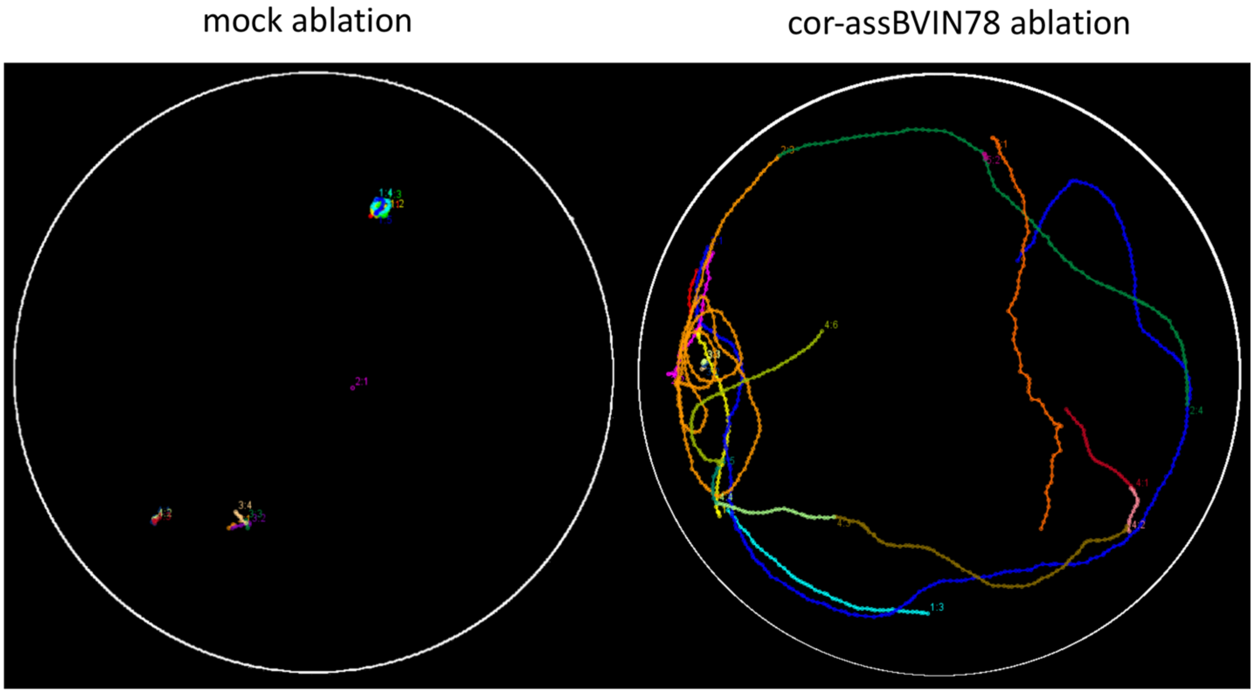


**Figure S5. Combined swim tracks from ablation Trial 2. Related to Figure 6.** The combined swim tracks of four mock ablated and five cor-assBVIN78 ablated larvae are shown. Each swim was given an arbitrary color.

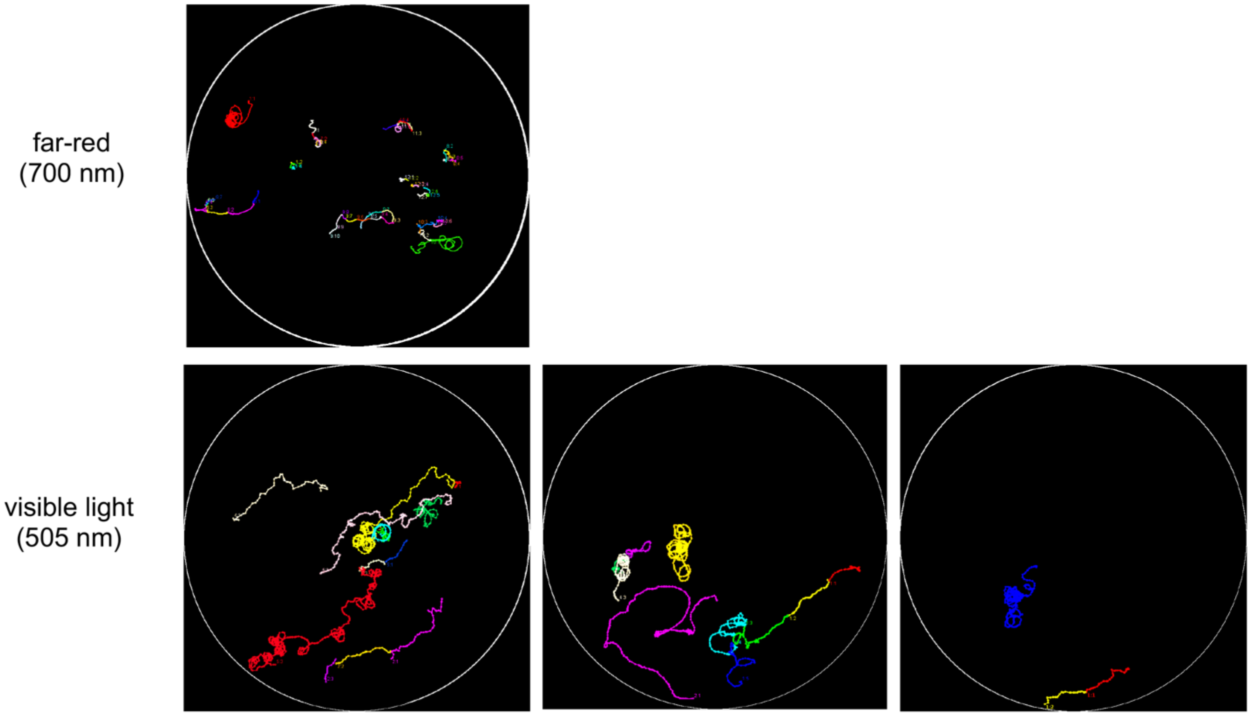


**Figure S6. Representative swim traces of unmanipulated larvae in the presence of directional visible light (505 nm), or far-red light (700 nm). Related to Figure 6.**
